# Latent learning drives sleep-dependent plasticity in distinct CA1 subpopulations

**DOI:** 10.1101/2020.02.27.967794

**Authors:** Wei Guo, Jie J. Zhang, Jonathan P. Newman, Matthew A. Wilson

## Abstract

Latent learning allows the brain the transform experiences into cognitive maps, a form of implicit memory, without reinforced training. Its mechanism is unclear. We tracked the internal states of the hippocampal neural ensembles and discovered that during latent learning of a spatial map, the state space evolved into a low-dimensional manifold that topologically resembled the physical environment. This process requires repeated experiences and sleep in-between. Further investigations revealed that a subset of hippocampal neurons, instead of rapidly forming place fields in a novel environment, remained weakly tuned but gradually developed correlated activity with other neurons. These ‘weakly spatial’ neurons bond activity of neurons with stronger spatial tuning, linking discrete place fields into a map that supports flexible navigation.

---

In his seminal 1948 paper, Edward Tolman proposed a ‘cognitive map’ theory, arguing that the brain organizes past experiences into a map-like memory that can serve as a generative guide for future actions ^1^. His argument gained a firm footing with the discovery of hippocampal place cells; their distinct spatial tuning functions offered a potential substrate to build mental maps of the environment ^2^. Recent studies showed that hippocampal neurons also encode a wide range of non-spatial information that enables cognitive functions beyond spatial navigation ^3–6^. However, we still lack a clear understanding of how cognitive maps emerge during learning ^2,7–9^. By studying subjects performing foraging or reinforcement learning tasks, we limited ourselves in observing neural activities associated with overt behavioral changes, whereas cognitive maps can be acquired and stored implicitly. For example, Tolman demonstrated a phenomenon known as latent learning, in which freely-exploring rats could covertly learn the layout of a maze without reinforced training or behavioral changes, suggesting that that internal state of the neural system is an essential aspect of memory ^1,10^. In this study, we investigated the latent learning of spatial maps in the hippocampal circuit. Using a manifold learning technique to visualize neuronal state space, we sought to explore the internal representation of spatial memory and further our understanding of the neural mechanism behind cognitive maps.

We expressed Ca^2+^ indicator jGCamp7f ^11^ in the dorsal hippocampus (HPC) of mice. With a miniature head-mounted single-photon microscope ^12^, we recorded Ca^2+^ activity of neurons in the pyramidal layer of the CA1 region through a chronically implanted GRIN lens. The animal was freely exploring in mazes of various geometric shapes with no reward or punishment association (Fig. 1A-C, Fig. S1). Neural data and the animal’s location were simultaneously recorded using the Bonsai software package ^13^. Raw Ca^2+^ recordings were analyzed with the CNMF-E algorithm in the CaImAn software package to isolate regions of interest (ROIs), which we treated putatively as neurons ^14^. Our recordings typically contained 50 – 300 neurons (Fig. 1D). We treated the population Ca^2+^ activity recorded during a session as an N-dimensional point cloud, where N is the number of neurons, and each point represents the ensemble neural state during a 100 ms window (Fig. 1E). Recent studies have shown that the intrinsic dynamics of a neural ensemble often occupy a low-dimensional manifold within the high-dimensional state space, which can be visualized after proper dimensionality reduction ^15–17^. We used Isomap, an unsupervised manifold learning algorithm, to find a low-dimensional embedding of the high-dimensional state space ^18^. The resulting manifold preserves the geodesic distance between all pairs of neural states if the transitions only consist of series of hops to nearby states, which mirrors the smooth state transitions in the HPC during spatial navigation induced theta oscillations ^19,20^. The parameters for the algorithm were chosen adaptively for each recordings session to minimize information loss and avoid overfitting (Fig. S2). Throughout this paper, we transformed all the recording sessions to 2D manifolds for easy comparison, although in many cases, extra dimensions provide information not contained in the first two (Fig. S3) ^17^.

**Fig. 1.**
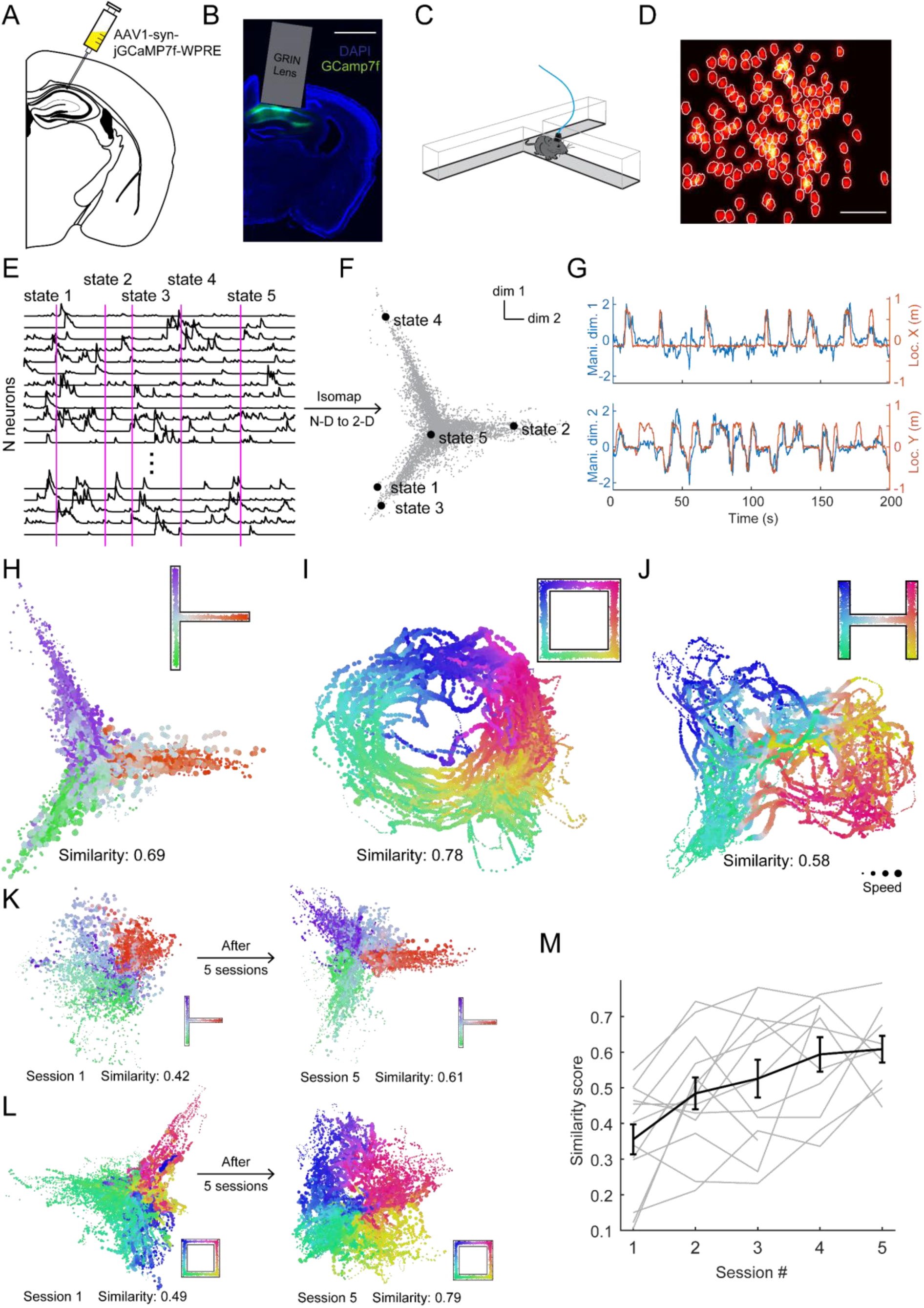
Latent learning induces plasticity in neural manifold representation of space in CA1. (A) Virus delivery of genes. (B) Coronal section of a mouse brain showing jGCamp7f expression (green) and GRIN lens location. Scale bar: 1mm. (C) Recording from a freely exploring mouse. (D) Example footprints of isolated ROIs. Scale bar: 0.1mm. (E) Example segment of Ca^2+^ traces of HPC neurons. (F) Example 2D manifold calculated from data in (E). The 2D states correspond to states labeled in (E). (G) Comparison of the neural state trajectory on the manifold and the animal’s spatial trajectory. (H-J) Example manifold representations of 3 mice running 3 different mazes. Each dot corresponds to a populational neural state during a 100 msec window, with the size of the dot representing the speed of the animal’s movement (range: 0-1 m/s), and the color of the dot representing the animal’s location in the maze (inset). (K-L) Example manifold representations of a maze early and late during latent learning. (M) Group data showing the improvement of similarity score throughout latent learning (F_4_ = 5.54, p = 0.002, repeated measures ANOVA. 13 mazes, 10 mice).

We observed that, if the animal was familiar with the maze, the animal’s spatial experience was encoded intrinsically in the neural manifold (Fig. 1F-G). Without information about the animal’s behavior or individual neuron’s spatial tuning, the algorithm uncovered a 2D manifold of the neural states that shared a close resemblance with the animal’s physical coverage in the maze (Fig. 1F, H). Moreover, the animal’s ‘mental trajectory,’ represented by the neural state transitions on the 2D manifold, reliably tracked the animal’s moment-to-moment physical trajectory in the maze (Fig. 1G). A similarity score between the mental and physical trajectories can be quantified based on their correlation coefficients. We observed high similarity scores in cases across multiple animals running in mazes with various shapes, suggesting faithful representations of spatial maps as the HPC internal states (Fig. 1H-J). Interestingly, the neural manifolds encoded the structures of the mazes topologically, turning a T maze into a Y and a square loop into a circle, which agrees with multiple works theorizing topological cognitive maps in the HPC ^21–23^.

However, if the animal was new to the maze, the neural manifold representation was less accurate (Fig. 1K-L). As the animal repeatedly explored the maze over a few sessions, the similarity score increased as the manifold looked more similar to the maze (Fig. 1K-M). This improvement was not accompanied by overt changes in the animal’s exploratory behavior, such as distance traveled, movement speed, or acceleration (Fig. S4). The intrinsic representation and the long-time-scale plasticity match the features of latent learning, suggesting the HPC neural manifold as one of the neural correlates of cognitive maps.

The similarity score between the neural manifold and the maze assesses the neural representation of the spatial map without resorting to explicit behavioral testing. This flexibility allows us to study the short-term effects of memory formation. For example, sleep is known to be involved in latent learning and memory consolidation ^24^. Does a single sleep session change the intrinsic representation of spatial maps? To test this, we exposed mice to a novel maze twice during the same day. Between the two runs, the animals spent 3 hours in their home cages (Fig. 2A-D). Half the mice (sleep group, Fig. 2A-B) were allowed to go to sleep, and the remaining (no-sleep group, Fig. 2C-D) were forced to stay awake (Fig. S5A-C). We compared the neural manifolds from the two maze sessions and found that the similarity score improved significantly if the animals were allowed to sleep, and showed no change if sleep was disrupted (Fig. 2E). These data suggest that changes in the HPC neural manifold are sleep-dependent.

**Fig. 2.**
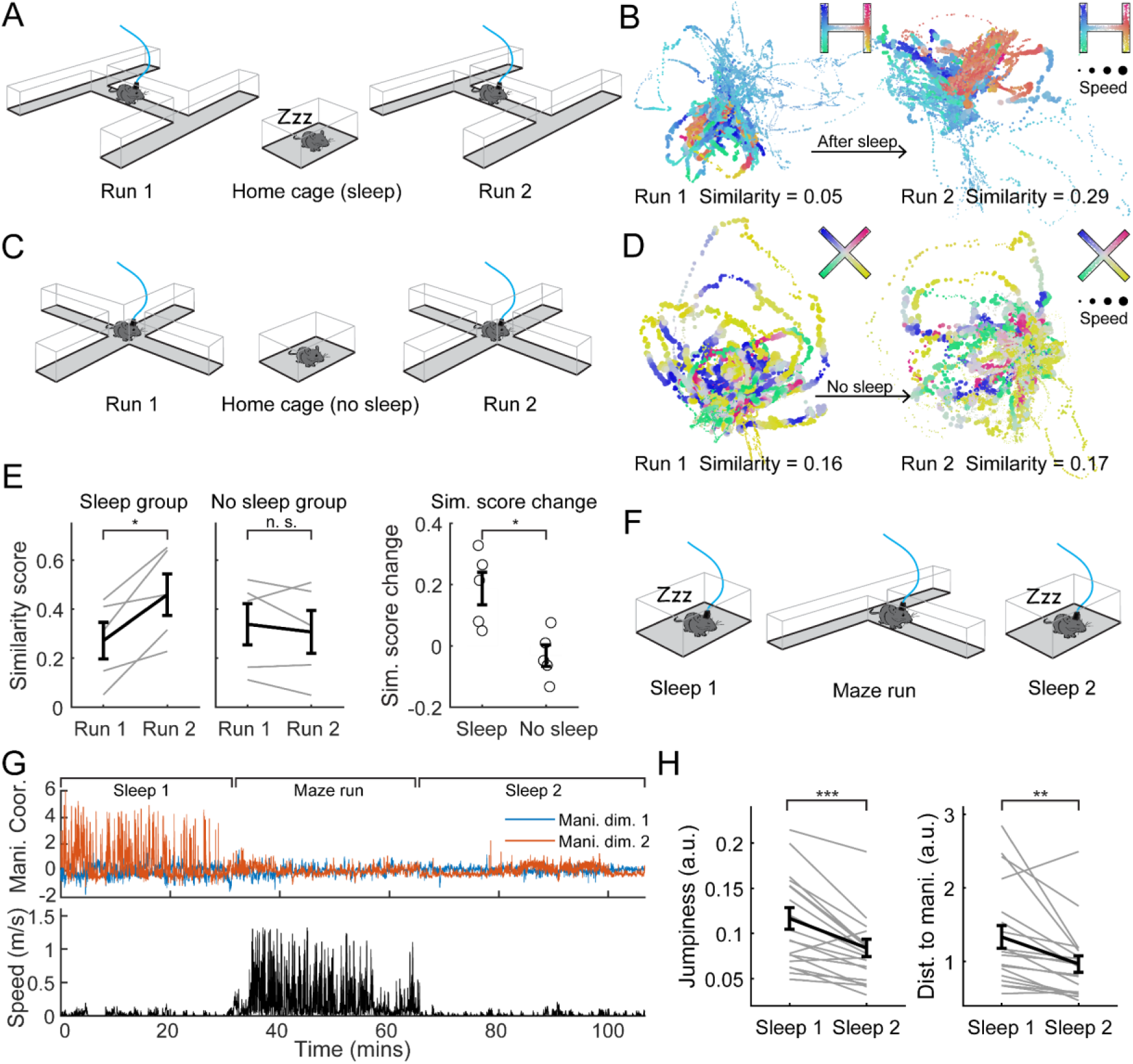
Neural manifold plasticity is sleep-dependent. (A, C) Cartoons showing the task. The mice are either allowed to or prevented from sleep during a 3 hour break between two maze runs. (B, D) Example neural manifolds from the sleep group (B) and the no-sleep group respectively (D). (E) Similarity score significantly improved in the sleep group (T_4_ = 3.51, P = 0.02, paired Student’s T-test, 5 sessions, 4 mice), but not in the no-sleep group (T_4_ = 0.9, P = 0.42, paired Student’s T-test, 5 sessions, 4 mice). There is a significant difference in the similarity score change between groups (T_4_ = 3.42, p = 0.009, unpaired Student’s T-test, 5 sessions, 4 mice). (F) Cartoon showing the task investigating neural activity during sleep before and after a maze run. (G) Top: Example trajectory of neural states on the manifold constructed from maze run data. Bottom: Animal’s speed. (H) Neural trajectory during the pre-run sleep is jumpier than the post-run sleep (T_18_ = 5.11, P = 7.25 × 10^−5^, paired Student’s T-test, 19 sessions, 4 mice); Neural states during the pre-sun sleep is further away from the manifold than the post-run sleep (T_18_ = 3.40, P = 0.008, paired Student’s T-test, 19 sessions, 4 mice). *: p < 0.05, **: p < 0.01, ***: p < 0.001.

Next, we investigated neural activity during sleep within the context of neural manifolds. We recorded the HPC activity of animals during a maze run session and two sleep sessions immediately before and after (Fig. 2F, Fig. S5D-E). We used the maze run session to establish a dimension reduction scheme, mapping the data to a 2D manifold. The same scheme was used to map the sleep data, effectively ‘decoding’ neural states during sleep as point clouds in the same subspace during the maze run (Fig. 2G). Comparing the two sleep sessions, we found that post- run sleep data were located closer to the manifold from the maze run and showed significantly less temporal ‘jumpiness’ (Fig. 2G-H). This suggests that during the post-run sleep, the HPC was reactivating neural states and sequences from the run, matching the well-described HPC replay phenomenon, which is essential for memory consolidation in the hippocampal-cortical circuit ^25,26^.

The HPC place code is known to form rapidly in a novel environment and remain stable afterward ^27–29^. However, the changing neural manifold demonstrates a slower consolidation of local circuit dynamics not described in previous studies. To investigate, we examined neurons’ spatial tuning across the latent learning sessions (data in Fig. 1M). For each neuron, we constructed two tuning functions during each maze run: one based on the animal’s physical trajectory (place field, Fig. 3A-B, top rows), and the other based on the mental trajectory (mental field, Fig. 3A-B, bottom rows). The selectivity of the tuning functions was calculated as their information content (I_p_ for place fields and I_m_ for mental fields) ^30^. As expected, a subset of HPC neurons immediately developed clear selectivity for physical locations after encountering a novel environment (high I_p_ values, Fig. 3A, right panels), while other neurons remain nonselective (low I_p_ values, Fig. 3A left panels). The mental fields and their information content I_m_ are similar to their place field counterparts, further demonstrating neural manifolds as spatial maps. After the animal gained more experience with the maze, both the I_p_ and I_m_ of the HPC population improved (Fig. 3B-C). We calculated a ratio between the I_m_ and I_p_ for each neuron and found that, on average, it grew monotonically over time, suggesting that I_m_ improved more than I_p_ during learning (Fig. 3D). However, not all neurons showed the same level of increase in I_m_. We found that in the last exploration sessions, high I_m_/I_p_ ratios were mostly found in neurons with low I_p_ values, i.e., neurons with weaker spatial selectivity (Fig. 3B, E). We performed the same analyses using the sparsity of spatial tuning as a measurement of selectivity ^30^ and reached the same conclusion (Fig. S6A-C).

**Fig. 3.**
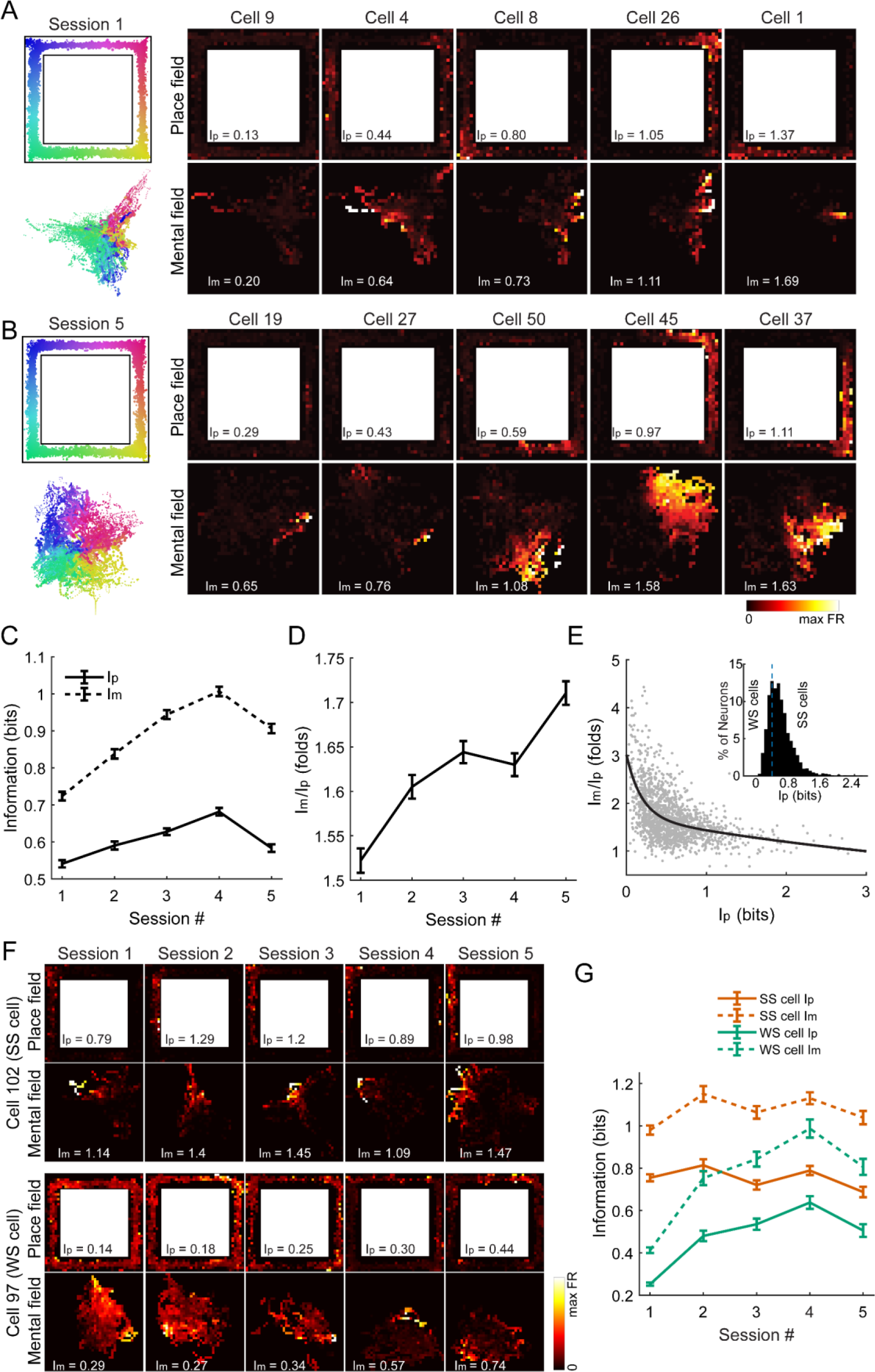
Latent learning induces heterogeneous neural tuning changes in CA1 neurons. (A) Example neurons’ place field tuning (top row) and mental field tuning (bottom row) from an early exploration session. Tuning quality ranges from low to high (left to right). (B) Similar to (A), but from a late exploration session. (C) Group data showing I_p_ and I_m_ changes over the course of learning (Main effect for time: F_4_ = 105.28, P = 8.38 × 10^−89^; main effect for measurement type (I_p_ versus I_m_): F_1_ = 336.4, P = 1.29 × 10^−309^; Two-way ANOVA. 1522 to 2076 neurons from 10 mice) (D) Group data showing changes of I_m_/I_p_ =over the course of learning (F_4_ = 27.1, P = 2.34 × 10^−22^; One-way ANOVA. 1522 to 2076 neurons from 10 mice) (E) High I_m_/I_p_ values are associated with low I_p_ values in the last session (WS cells versus SS cells: F_1_ = 323.37, P = 1.06 × 10^−63^; One-way ANOVA. 1522 neurons from 10 mice). Inset: distribution of I_p_ during during the last session. Mode: 0.4. (F) Example tuning functions of tracked neurons over the course of learning. Top: place cell. Bottom: non-place cell. (G) Information change of tracked place and non-place cells over learning (SS cells: main effect for time: F_4_ = 0.7, P = 0.58; main effect for measurement type: F_1_ = 172.0, p = 9.49 × 10^−34^, interaction: F = 1.37, P = 0.24. WS cells: main effect for time: F_4_ = 1.4, P = 0.25; main effect for measurement type: F_1_ = 92.5, p = 1.05 × 10^−18^, interaction: F = 3.1, P = 0.01. 330 SS cells, 176 WS cells from 10 mice. Repeated measures ANOVA).

Most studies of the HPC circuit have focused on neurons with clear spatial tuning. However, recent studies suggested that neurons with individually weak spatial selectivity can carry significant spatial information as an ensemble ^31^. Were these weakly-tuned neurons responsible for shaping the neural manifold? To investigate, we used the CellReg algorithm ^32^ to identify neurons we consistently recorded across all sessions (Fig. 3F) and divided them into two groups based on their initial I_p_: strongly spatial cells (SS cells, I_p_ > 0.4 bit) and weakly spatial cells (WS cells, I_p_ <= 0.4 bit), where the threshold was chosen to be the mode of the I_p_ distribution (Fig. 3E inset). We found that the original SS cells remained stable and showed relatively flat I_p_ and I_m_ growth over time, while the original WS cells underwent striking changes in both place and mental fields and significant improvement in both I_p_ and I_m_ (Fig. 3F-G). These data suggest that the changes in the WS cells may be responsible for changes in the neural manifold during latent learning.

Conceptually, neural state space takes the form of a low-dimensional manifold when there is redundancy in the neural code, which often leads to correlated activity among neurons (Fig. 4A) ^17,22^. Identifying changes in neural correlation may reveal the mechanism behind the manifold plasticity. For each neuron, we defined a correlation score as its mean absolute correlation with all other neurons. During latent learning, the population correlation score increased modestly (Fig. 4B). However, once neurons were divided into SS and WS cells based on their spatial tuning (threshold = 0.4 bit), we discovered that the main source of correlation increase was the WS cells (Fig. 4B), which became much more correlated with both SS cells and other WS cells during latent learning (Fig. S6E). This result explains and linked two earlier observations. First, as a WS cell became more correlated with other neurons, it developed higher selectivity within the neural state space, resulting in higher I_m_ in the mental field tuning. Second, elevated correlation of WS cells with SS cells introduces redundancy in the ensemble tuning, turning the neural state space into a low-dimensional manifold.

**Fig. 4.**
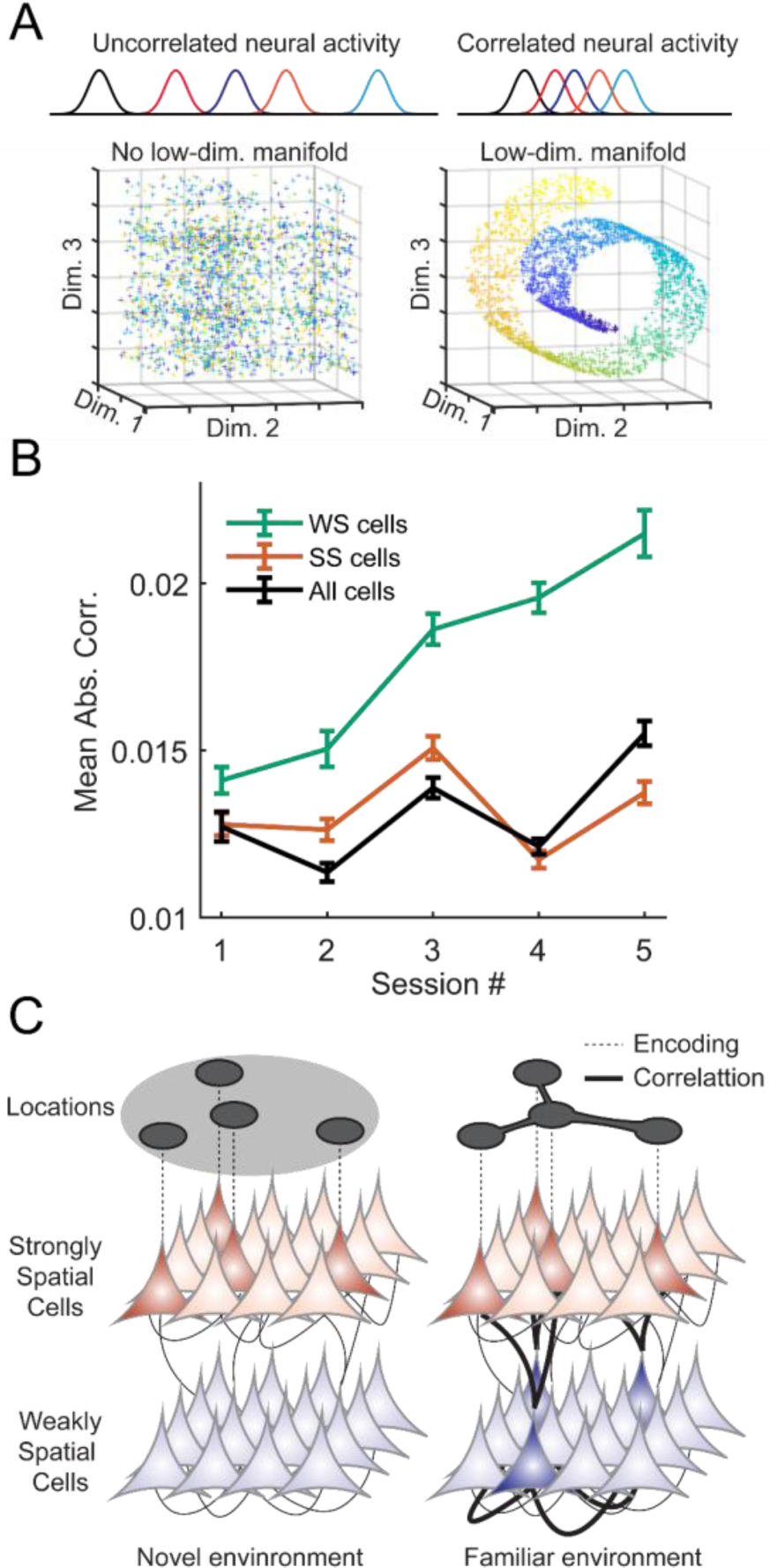
Latent learning elevates neural correlation in weakly spatial cells. (A) Cartoons showing relationship between neuronal correlation and neural manifolds. (B) Correlation score change over learning. (Main effect for time: F_4_ = 20.6, P = 6.48 × 10^−17^, main effect for cell type: F_1_ = 290.55, P = 1.81 × 10^−64^, two-way ANOVA. 330 SS cells, 176 WS cells from 10 mice) (C) Cartoon showing the conceptual model for latent learning.

To summarize our findings, we propose the following as a mechanism for the latent learning of cognitive maps. When the animal first encounters a new environment, a subset of neurons in the HPC quickly develop preferential tuning for discrete locations, becoming place cells (SS cells). Later, ensemble cell activities representing the spatial experience are reactivated in the HPC during sleep. With repeated exposures to the environment and sleep afterward, the previously non-selective neurons (WS cells) start to develop correlated firing activity with specific SS cells, and with each other. The changes in WS cells add redundancy to the population code, turning the neural state space into a smooth low-dimensional manifold. As a result, the WS cells form connections that topologically link discrete place fields of the SS cells into a map structure (Fig. 4C).

Why does the brain make an effort to turn an already accurate place code into a map? As Tolman argued, maps are efficient, flexible, and most importantly, generative. Indeed, numerous works have demonstrated that the HPC circuit is not merely a location recognizer, but a trajectory generator. Phenomena such as theta sequences during navigation ^19,20^ and awake replays during the task solving process ^25^ all rely on a map-like neural code in the HPC. We provided a mechanism showing how a cognitive map can be built via latent learning and stored as a neural manifold. Although we only investigated simple 2D spatial exploration experiences, with our framework, non-spatial information such as sensory cues, reward, behavioral states, and even the concept of time can all be stored in same the manifold, likely in the other dimensions, providing a holistic, schema like representation of the animal’s experience ^2,7–9,33^. Our work also showcased the often- neglected WS cells in the HPC, demonstrating the importance of understanding heterogeneity in the HPC neurons ^34,35^. Previous work has shown that neurons in different sublayers of CA1 exhibit different levels of stability, which may be contributing to the plasticity of different time scales we observed ^36^. In sum, our findings provide a novel approach to study the HPC circuit, leading to new insights into the mechanisms of learning and memory.

## Acknowledgments

We thank Daniel Cho and Qianli Xu for assisting the experiments. We thank Dr. Daniel Polley for reading and commenting on the manuscript.

## Author contributions

WG and MAW designed the experiments; WG and JJZ collected the data with techniques developed by JPN; WG performed the analysis and prepared the manuscript; MAW supervised the entire project.

## Data and materials availability

Data and analysis code can be made available upon request to any researcher for purposes of reproducing or extending the analysis.

## Materials and Methods

### Subjects

All procedures were approved by the Massachusetts Institute of Technology Committee on Animal Care and followed the guidelines established by the National Institutes of Health for the care and use of laboratory animals. A total of 16 male mice of either C57/B6 or CBA/CaJ background were used in the experiments.

### Virus-mediated gene delivery

Mice were initially anesthetized with 5% Isoflurane in oxygen and moved to a stereotaxic surgical frame (Kopf). Throughout the procedure, the animals were maintained at a surgical plane of anesthesia using continuous 1.5% to 2% Isoflurane in oxygen, with the body temperature maintained around 36.5°C using a homeothermic blanket system (Kent Scientific). We numbed the surgical area with a subcutaneous injection of lidocaine (5 mg/mL), then made an incision along the midline of the scalp to expose the skull. A burr hole was made on the skull with a dental drill -1.5mm medial-lateral and 2.1mm rostral-caudal to the Bregma landmark. Using a motorized injector, we injected 0.5µL of pGP-AAV1-syn-jGCaMP7f-WPRE (titer around 1.9 x 10^12^ gc/ml) in the burr hole 1.5mm under the skull surface at a speed of 0.05 µL/min. After the end of the injection, the pipette was left in the brain for 5 minutes before pulling out. Following the procedure, the scalp was sutured back, and analgesic was administered (Buprenex, 0.05 mg/kg) prior to animals’ recovery.

### GRIN lens implant

One to two weeks after virus injection, mice were once again brought to a surgical plane of anesthesia using the same protocol for general anesthesia, local anesthesia, and body temperature control described above. First, the scalp was removed with the periosteum overlying the dorsal surface of the skull thoroughly cleaned with 70% ethanol. The skull surface was roughened with an etchant gel (C&B Metabond), then thoroughly cleaned once again. A craniotomy 2 mm to 2.2 mm in diameter was made on the skull, containing the original injection burr hole in the center. Cortical tissue was carefully aspirated out, exposing the rostral-caudal running striation pattern right above dorsal CA1. A GRIN lens (1.8 DIA, 0.25 pitch, 670nm, Edmund) was carefully placed on top of CA1 and fixed to the skull with adhesive (Loctite 454). Lastly, we covered the entire exposed skull and the side of the GRIN lens first with clear dental cement (C&B Metabond), then with black cement (fast curing orthodontic acrylic resin, Lang) to block out ambient light. To protect the GRIN lens, we covered the exposed surface with fast curing silicone gel (KwikSil). After administering analgesic (Buprenex, 0.05 mg/kg), animals were woken up and given at least two weeks of recovery time.

### Baseplate implant

Mice were once again anesthetized similar as described above. With a baseplate attached to a miniscope (UCLA miniscope) ^12^, we image through the GRIN lens and manually adjust the angle and the focus until an optimal field of view and image quality were reached. The baseplate was first fixed in place with adhesive (Loctite 454), then securely affixed to the skull with dental cement (M&B metabond). The animals were allowed to recover afterward.

### Calcium imaging in freely moving animals

Once the animals were thoroughly recovered from previous procedures, we start in vivo awake recording sessions. First, animals were allowed to acclimate to the miniscope and tethering for a few sessions in the home cage. We also use this opportunity to apply fine adjustments of miniscope focus and illumination power to achieve optimal image quality. Once animals exhibit normal behavior and move without trouble, we proceed to maze exploration experiments. The enclosures were made out of corrugated plastic sheets (Home Depot), and their shapes and dimensions described in Fig. S1. We used software package Bonsai to record Ca^2+^ activity from the miniscope (30Hz sampling rate) and the animal’s behavior using a webcam (60Hz sampling rate) simultaneously (https://github.com/jonnew/Bonsai.Miniscope). The animal’s location was identified by tracking a red tape attached to the head-mounted miniscope against the white floor background.

### Sleep recording and sleep disruptions

For Ca^2+^ recordings during sleep or quiet wakefulness, miniscope tethered animals were allowed to move freely in their homecage located in an enclosure with a different visual context from the mazes (Fig. S1). Recording starts once the animal exhibits prolonged immobility and lasts around 30 mins. The animal’s movement was continuously monitored during recording. For maze running recordings after sleep, we wait until the animal fully wakes up and shows normal exploratory behavior.

For sleep disruption experiments, the animals were placed in their homecage for 3 hours between two maze runs. Mice in the sleep group were left undisturbed for 3 hours. Mice in the no-sleep group were prevented from going to sleep by the experimenter infrequently tapping on the cage or rustling the bedding, which also lasted for 3 hours. The animals’ movement was tracked using real-time computer vision algorithms (CV package in Bonsai). We putatively define the animal’s state as asleep of quiet wakefulness when the speed of the locomotion is below 5 cm/s for over 0.5 minutes.

### Ca^2+^ Data Analysis

The Ca^2+^ recordings were analyzed with the software package CaImAn using the CNMF-E algorithm ^14^. Parameters for each animal’s recording were optimized to ensure optimal extraction results. The first and last 1000 frames of the recordings were not included in the analysis. We used the denoised Ca^2+^ dynamics (variable C in the CNMF-E model) as the ROIs’ activity time series. ROIs with weak (peak C value lower than 100% df/f) or sporadic activity (less than three instances of activity higher than 100% df/f) were eliminated from further analysis. The selected ROIs were putatively treated as neurons.

### Manifold discovery

The recording was first downsampled from 30Hz to 10Hz. Each neuron’s Ca^2+^ response was normalized between the values 0 and 1. This resulted in an N x T matrix where N was the number of neurons, and T was the number of frames in the recording. We used the Isomap algorithm (scipy-learn) to embed the neural data to 2D (3D in Fig. S3), which yielded a 2 x T matrix. To find an optimal embedding of neural recordings, we try a series of numbers of nearest neighbors (NNs) for the Isomap algorithm range from 5 to 100 in steps of 5. Each time we increased the NNs, we compare the reconstruction error of the new embedding against the previous one. Once the improvement of reconstruction error plateaus (less than 10% increment for two NNs in a row), the algorithm stops. Empirically we found that manifolds discovered with this approach share the closest resemblance to the animal’s spatial experience without starting to breakdown due to overfitting (Fig. S2E-H).

To calculate the similarity score between a neural manifold and the enclosure, we first need to find an angular rotation of the manifold that best matches the maze. We rotated the neural manifold with respect to the origin point in 5-degree increments (Fig. S2A-D). For each rotated manifold, we calculated two correlation coefficients between the manifold trajectory and the animal’s physical trajectory (manifold dim. 1 versus the x coordinate; manifold dim. 2 versus the y coordinate). The average of the two is the similarity score between the manifold and the maze. Out of all the rotated manifolds, the one with the highest similarity score was chosen as the best-matched manifold and its similarity score as the similarity score of the manifold.

For the sleep data analysis, the neural manifold was calculated similarly using only data during the maze run portion. The same mapping was used to embed recordings during the sleep sessions to the neural subspace during the run. As a result, the high-dimensional data during sleep were mapped to a series of 2D points. The jumpiness of the resulting data was calculated as the absolute value of temporal changes in coordinates averaged over time, then averaged over the two dimensions. The distance to the manifold was first calculated for individual neural states as the average distance to all the points in the manifold during the maze run; then, the values were averaged across all neural states during the sleep session.

### Place field and mental field calculation

To calculate the place field of a neuron, the entire field of view, including the parts the animals could not travel to, was divided into a 40×40 grid. For each grid, we calculate the average neural response whenever the animal traverses the location as the place field response. As per convention, neural responses of the animal during fidgeting or quiet wakefulness (speed < 10 cm/s) were not used in place field calculation. The mental field of the neuron was calculated the same way except for using the 2D neural manifold space, which was also divided into a 40×40 grid. The information content of a neuron’s place field or mental field tuning was calculated as:

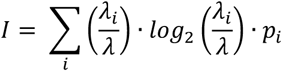

And the sparsity is calculated as:

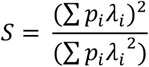

Where p_i_ is the probability of the animal occupying the *i*th bin (occupancy time divided by total time), λ_i_ is the average neural response in the *i*th bin, and λ is the overall average neural response. Because we used normalized neural activity in this calculation, the unit of the information content was simply referred to as bits.

We putatively divide neurons into strongly spatial (SS) cells and weakly spatial (WS) cells based on their place field information content. The dividing threshold is chosen to be the mode of the distribution (0.4 bits).

### Neural correlation analysis

A neuron’s correlation score was calculated as the mean absolute value of the correlation coefficients between its activity and all other neurons activity. This measure can be restricted to calculating correlation with only SS cells or WS cells.

### Cell identity tracking over sessions

For recordings obtained within the same day, we concatenated multiple blocks of the data used the CNMF- E algorithm to obtain consistent cell identities. For recordings obtained over multiple days, individual CNMF-E analysis was applied to separate sessions, and the CellReg algorithm was used to perform alignment based on the spatial correlations of the ROI footprints (variable A in the CNMF-E model). Only cells consistently showed up across all learning sessions were used in the analysis of Fig.3.

### Histology

Animals were deeply anesthetized and sacrificed with fatal plus (100mg/kg). The animal was first perfused with saline, then fixed with 10% formalin solution. The brain was removed and post-fixed in 10% formalin solution overnight before transferred to 30% sucrose solution in 0.01M phosphate-buffered saline. Coronal sections (50 µm thick) were cut with a cryostat (Leica), counterstained with DAPI (Vectashield), and photographed under a fluorescent microscope (Nikon Ti).

### Statistical analysis

All statistical analysis was performed with MATLAB (Mathworks). Descriptive statistics are reported as mean ± SEM unless otherwise indicated. Lillefors test was performed confirm normality of the dataset if parametric statistical tests (student’s T tests, ANOVA, etc.) were used. Non-parametric statistical tests were used in select cases where data samples were not normally distributed. In cases where the same data sample was used for multiple comparisons, we used the Holm-Bonferroni correction to adjust for the increased probability of Type-I error. Statistical significance was defined as p < 0.05.

---

**Fig. S1.**
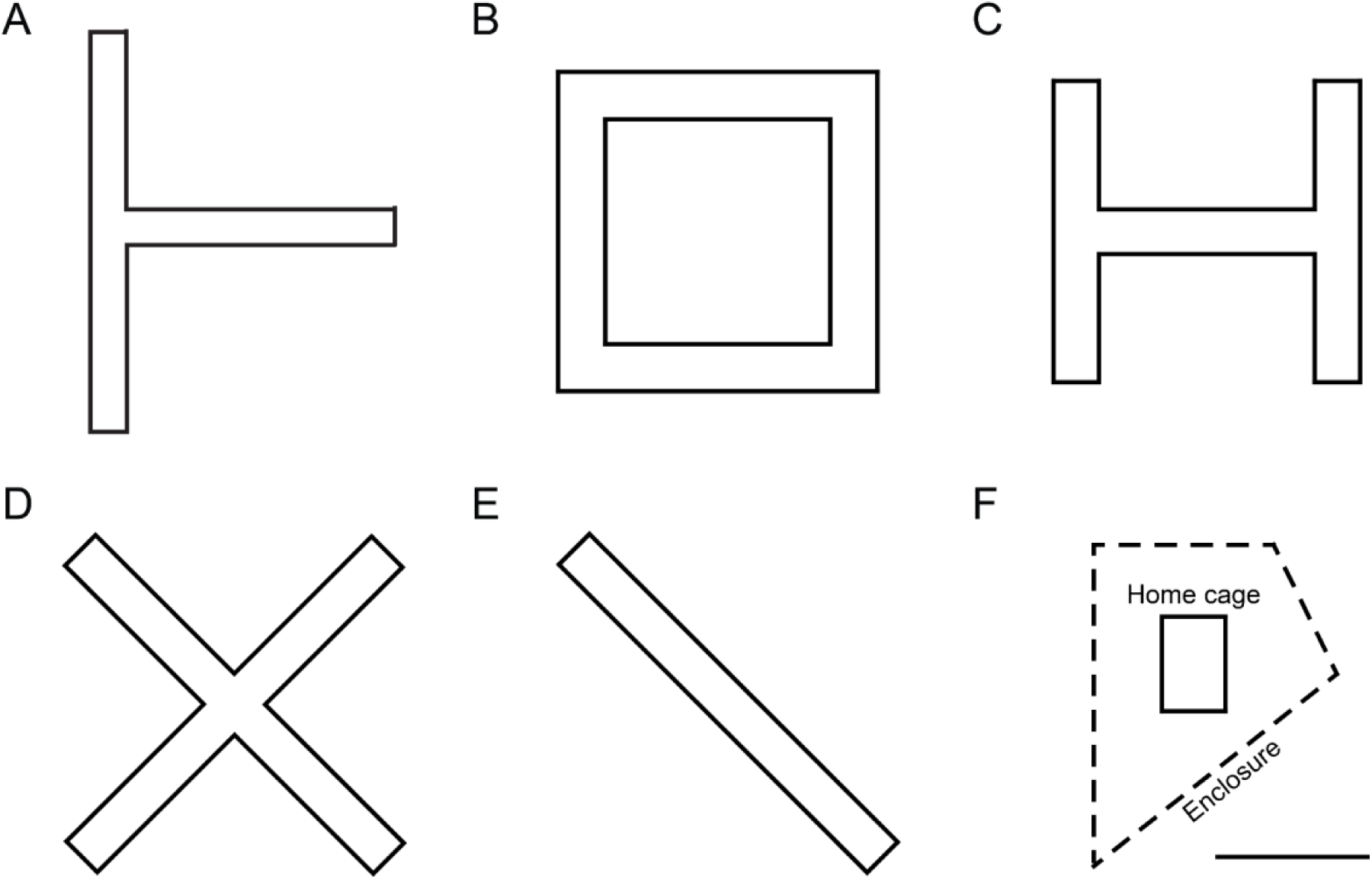
Geometric shapes of mazes used in the experiments. (A-E) Mazes. (F) Home cage placement. Scale bar: 0.5m.

**Fig. S2.**
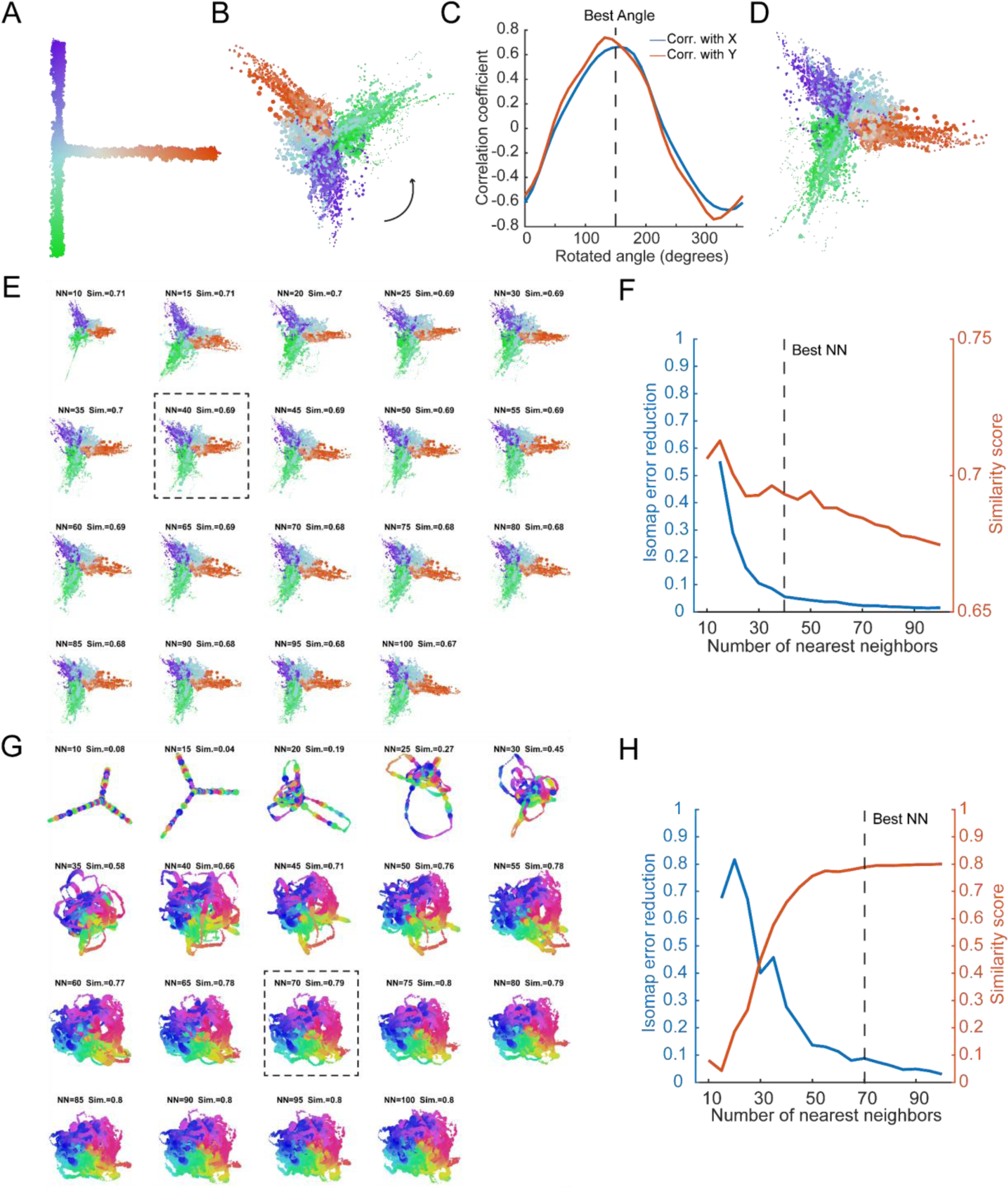
Parameter optimization for manifold discovery. (A-D) Optimal angle for manifold-maze alignment. (E-F) Two examples of identifying the optimal NN for manifold discovery. Using the criterion (less than 10% reduction in embedding construction error), we can identify the manifold that optimally matches the maze (plateau in the similarity score curve, boxed manifold).

**Fig. S3.**
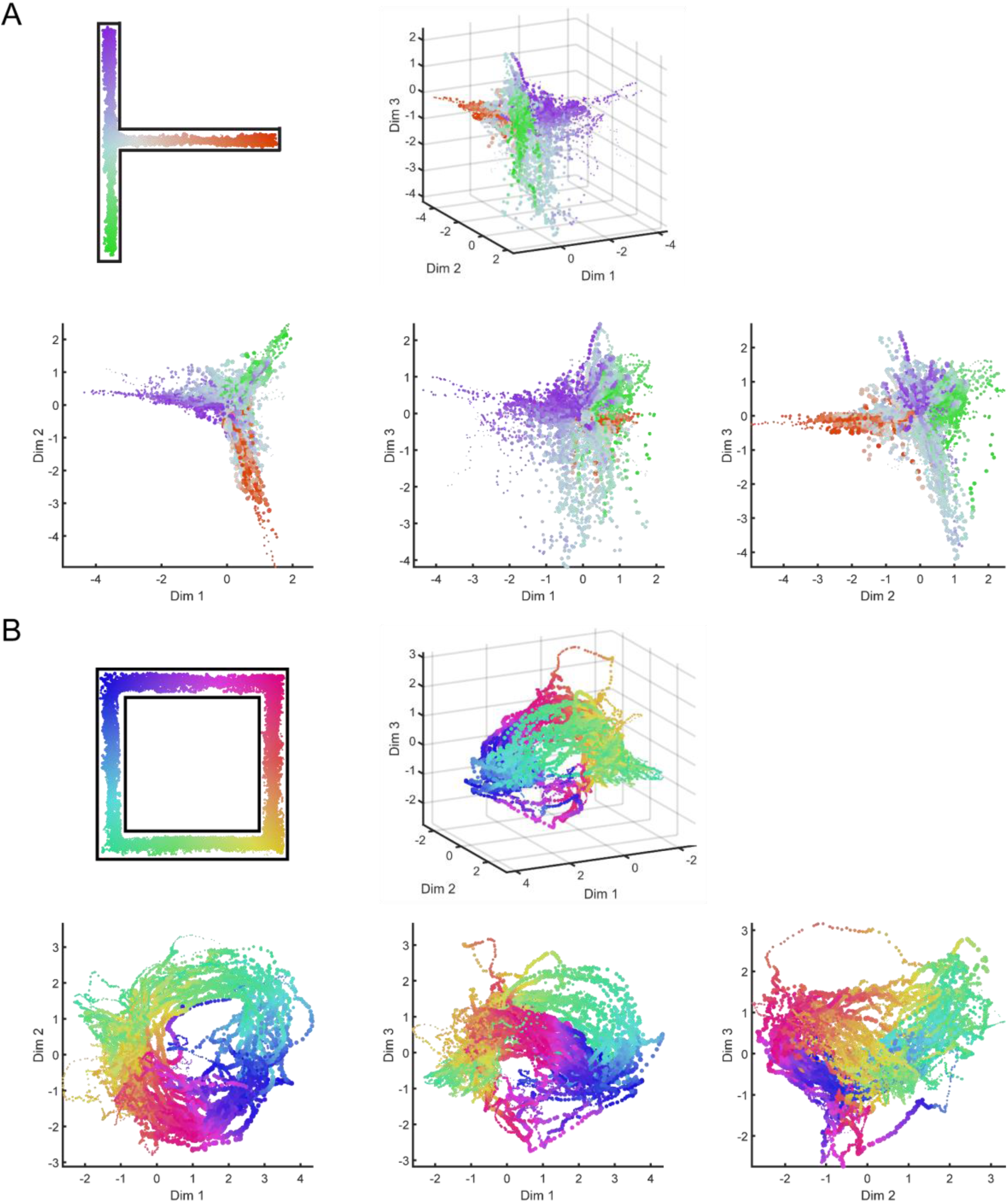
Extra dimensions of the neural manifold. (A) 3D manifold representation of data in Fig 1H. (B) 3D manifold representation of data in Fig 1I.

**Fig. S4.**
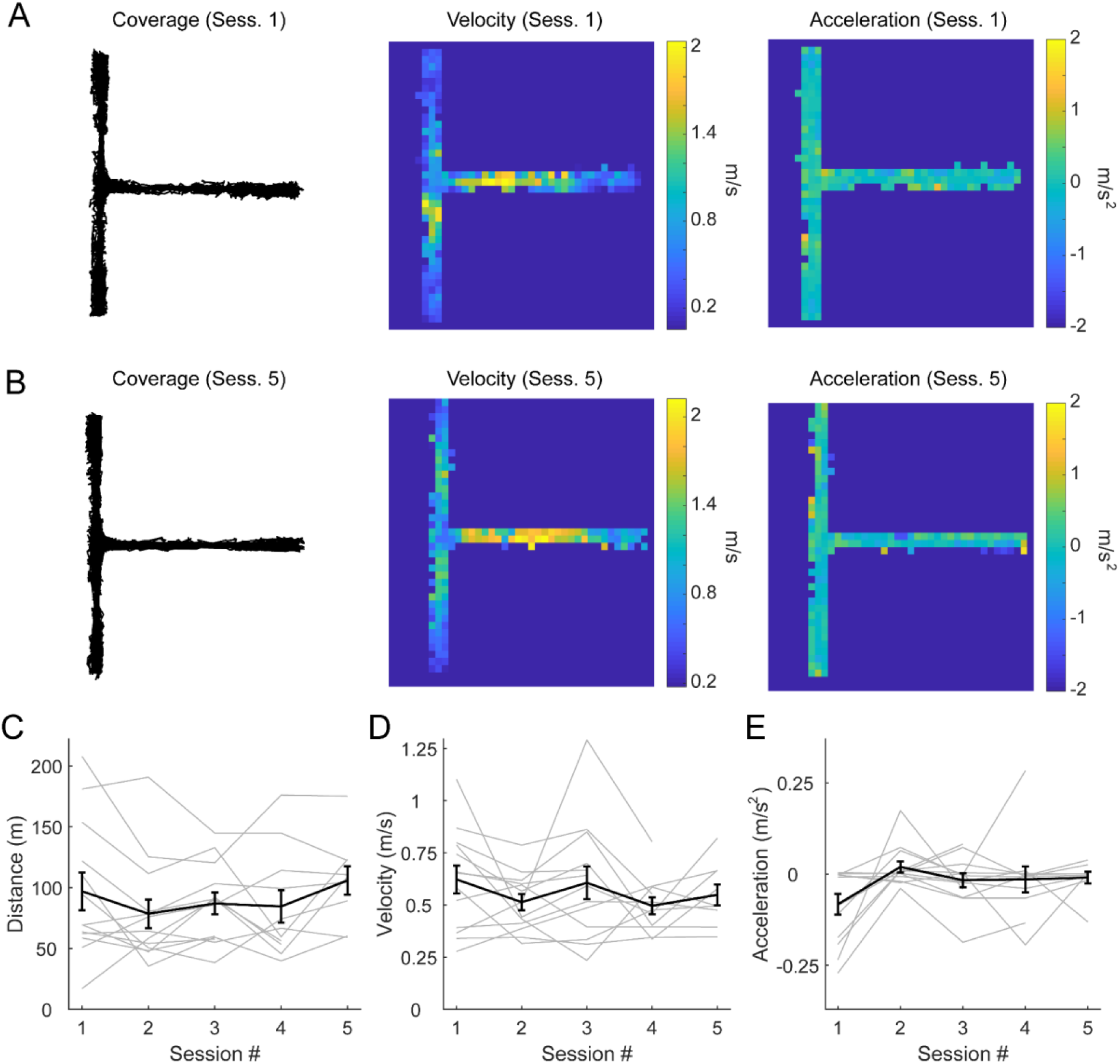
Overt behavior of animals during latent learning does not change. (A) Example coverage, velocity, and acceleration of mouse’s first session exploring a maze. (B) Data from the same mouse in (A) during its fifth session in the same maze. (C) Average distance travelled of the animals during exploration (Main effect for time: F_4_ = 1.79, P = 0.15, repeated measures ANOVA. 13 mazes, 10 mice). (D) Average velocity of the animals during exploration (Main effect for time: F_4_ = 1.53, P = 0.21, repeated measures ANOVA. 13 mazes, 10 mice). (E) Average acceleration of the animals during exploration (Main effect for time: F_4_ = 0.94, P = 0.71, repeated measures ANOVA. 13 mazes, 10 mice).

**Fig. S5.**
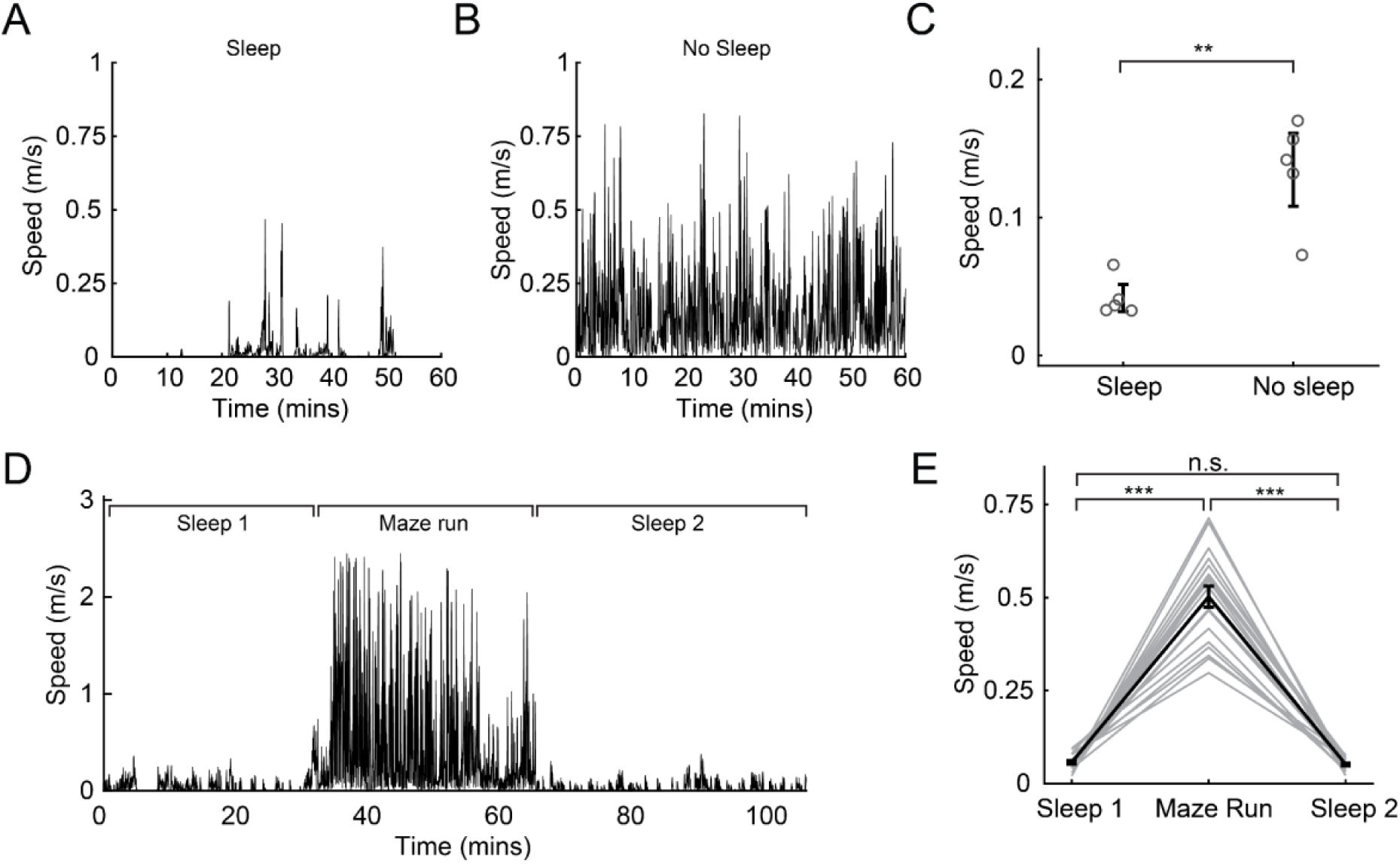
Movement tracking during sleep. (A) Example trace of animal’s speed from the sleep group. (B) Example trace of animal’s speed from the no-sleep group. (C) Group data showing the average speed during home cage stay was significantly different between groups (T_4_ = 6.1, P = 0.004, unpaired Student’s T-test. 5 sessions, 4 mice). (D) Example trace of an animal’s during the sleep-run-sleep. (E) Group data showing the average speed of animals during the 3 stages of the session (Sleep 1 versus Run: T_18_ = 14.3, P = 2.91 × 10^−11^; Sleep 2 versus Run: T_18_ = 15.4, P = 7.90 × 10^−12^; Sleep 1 versus Sleep 2: T_18_ = 0.8, P = 0.43. With Holm-Bonferroni correction for multiple comparisons. 19 sessions, 4 mice).

**Fig. S6.**
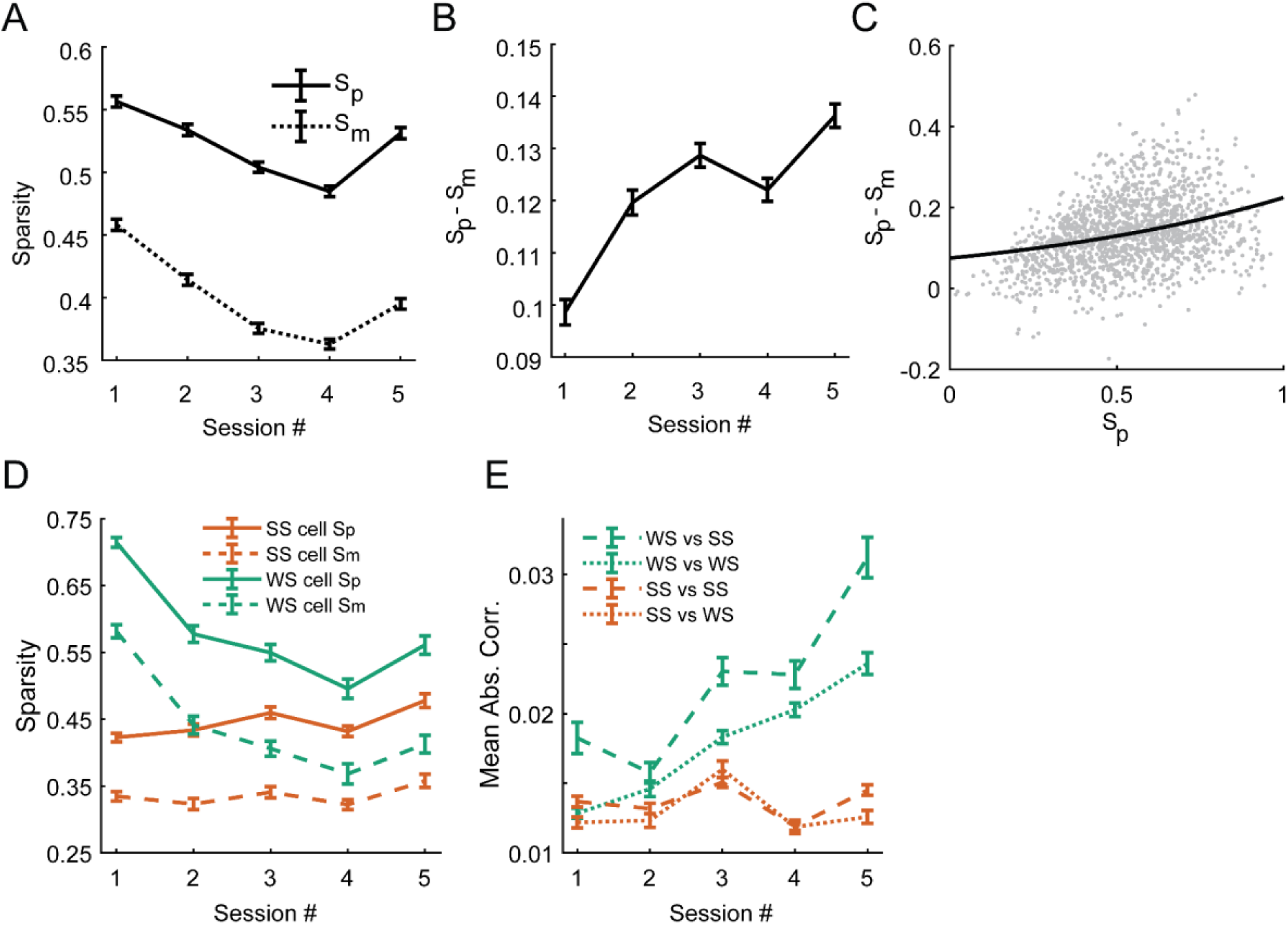
Additional tuning analysis for SS and WS cells. (A) Sparsity analysis of HPC neurons during latent learning (S_p_: place field sparsity, S_m_: mental field sparsity). (Main effect for time: F_4_ = 120.28, P < 2.23 × 10^−308^; main effect for measurement type (S_p_ versus S_m_): F_1_ = 1940.05, P < 2.23 × 10^−308^; Two-way ANOVA.1522 to 2076 neurons from 10 mice.) (B) Change of S_p_-S_m_ difference during latent learning. (F_4_ = 38.5, P = 5.5 × 10^−32^; One-way ANOVA.1522 to 2076 neurons from 10 mice.) (C) Distribution of S_p_-S_m_ as a function of S_p_ during late sessions (WS cells versus SS cells: F_1_ = 188.2, P = 1.80 × 10^−40^; One-way ANOVA. 1522 neurons from 10 mice). (D) Sparsity changes of tracked neurons. (SS cells: main effect for time: F_4_ = 2.16, P = 0.07; main effect for measurement type: F_1_ = 170.0, p = 2.02 × 10^−33^, interaction: F = 0.93, P = 0.44. WS cells: main effect for time: F_4_ = 10.5, P = 2.81 × 10^−8^; main effect for measurement type: F_1_ = 107.5, p = P = 4.9 × 10^−21^, interaction: F = 0.2, P = 0.96. 330 SS cells and 176 WS cells from 10 mice) (E) Correlation score changes of neurons, split into two groups: SS cells and WS cells. (WS versus SS: F_4_ = 60.0, P = 9.95 × 10^−50^; WS versus WS: F_4_ = 27.9, P = 7.91 × 10^−23^; SS versus SS: F_4_ = 12.82, P = 2.16 × 10^−0^; SS versus WP: F_4_ = 11.04, P = 6.84 × 10^−9^; All main effects for time, one-way ANOVA. 330 SS cells and 176 WS cells from 10 mice).

